# μGrowthDB: querying, visualizing, and sharing microbial growth curve data

**DOI:** 10.1101/2025.10.13.682118

**Authors:** Andrey Radev, Julia Casado Gómez-Pallete, Sofia Monsalve Duarte, Karoline Faust, Haris Zafeiropoulos

## Abstract

Microbial growth curve data contain valuable information about microorganisms’ physiology and metabolism. However, they are currently reported in various formats scattered across publications. A generic resource that allows querying and comparing microbial growth curve data is currently missing. For this reason, we developed μGrowthDB, an open repository designed to store, visualize, analyze, and share quantitative measurements of species abundances and metabolite profiles from monocultures and communities of known composition. μGrowthDB is designed to accommodate a range of experimental designs and measurement techniques. In addition to querying microbial growth curves by organism or metabolite, μGrowthDB supports comparative visualisation of growth curves and offers a guided data submission process intended to make data upload as easy as possible. μGrowthDB is available at: https://mgrowthdb.gbiomed.kuleuven.be/.

## Introduction

Microbial growth curves record the abundance of one or several microorganisms in a specific environment over time and often include additional measurements such as pH or metabolite concentrations. Microbial growth curves are collected for a number of reasons, e.g., to quantify minimal inhibitory concentrations (MIC) of antibiotics (Balouiri et al., 2016), to estimate food shelf-life (Labuza & Fu, 1993), or to improve wastewater treatment (Li et al., 2025; Vital et al., 2010). In general, they are also often required when developing and parameterizing quantitative models of microbial communities (Baig et al., 2023; Ram et al., 2019; Sprouffske & Wagner, 2016), which in turn help design communities and interventions to improve outcomes in applications ranging from healthcare to agriculture (Clark et al., 2021; Hromada & Venturelli, 2023; Xu et al., 2019). Despite their importance, there is currently no resource that allows structured storage and/or querying of existing microbial growth curve data. In the literature, growth curves are often only shown in plots or hidden in supplementary material. Thus, growth curve data are currently not easily *Findable* or *Accessible*, nor is there a standard format that would make them *Interoperable* (i.e., easy to combine with other data). Since they are scattered across research articles, existing growth curve data are also not easily *Reusable*, e.g., for the design of new experiments. In brief, there is not yet a FAIR resource available for microbial growth curve data. In food microbiology, ComBase (Baranyi & Tamplin, 2004) allows sharing and visualizing growth curve data, but it is specific to spoilage organisms and is not designed for microbial community data. This lack of a generic and FAIR resource of microbial growth curve data motivated the work presented here.

For a database of microbial growth curves to be useful, several challenges need to be overcome: the database has to accommodate heterogeneous experimental designs and measurement techniques, it needs flexible visualisation tools that allow comparing growth curves across conditions, and it has to make data upload as easy as possible. Growth curve data are too complex to be parsed in bulk from existing sources by a single team, and thus, a well-designed data submission process is essential to populate the database through a community effort.

Here, we present μGrowthDB, the microbial (μ) growth curve database, which addresses these challenges and is available at https://mgrowthdb.gbiomed.kuleuven.be/. In brief, μGrowthDB provides structured data storage and supports user-friendly uploading, visualization, analysis, and sharing of abundance and metabolites measurements for one or several microbial strains.

## Materials and Methods

### Database Schema

The database schema was developed in discussion with microbiologists to accommodate a wide range of experimental designs and measurement techniques. For this, the schema uses the following key concepts:

- A *Biological replicate* represents the culture unit in which one or more microbial strains were grown, e.g., a well, a bottle, a bioreactor, a plate, or complex configurations connecting more than one vessel (e.g.,(Van de Wiele et al., 2015))
- The distinct sections or partitions within a biological replicate are represented as *Compartments*. By default, a biological replicate consists of one compartment, but it may be linked to more than one to account for semipermeable membranes, beads, or other sub-structures inside the vessel or for several connected vessels.
- A *Community* is the set of strains growing in a biological replicate.

Growth curve data are then organized into three hierarchical levels: *Experiments, Studies*, and *Projects*, with a unique identifier each (see Figure 1). *Experiments* are defined as the lowest-level data entity, combining *Compartments* with *Communities* and a set of *Biological Replicates*; multiple *Compartments* and/or *Communities* may be included in an *Experiment* to describe experimental designs involving perturbations. *Experiments* belong to a *Study*, defined as a set of *Experiments* addressing a common hypothesis or question. A *Study* may consist of more than one *Experiment* (e.g., a mono- and a co-culture, each with several biological replicates), and it can be annotated with a publication URL. *Studies*, in turn, belong to *Projects*, which group studies addressing a specific research question or topic.

**Figure 1.**
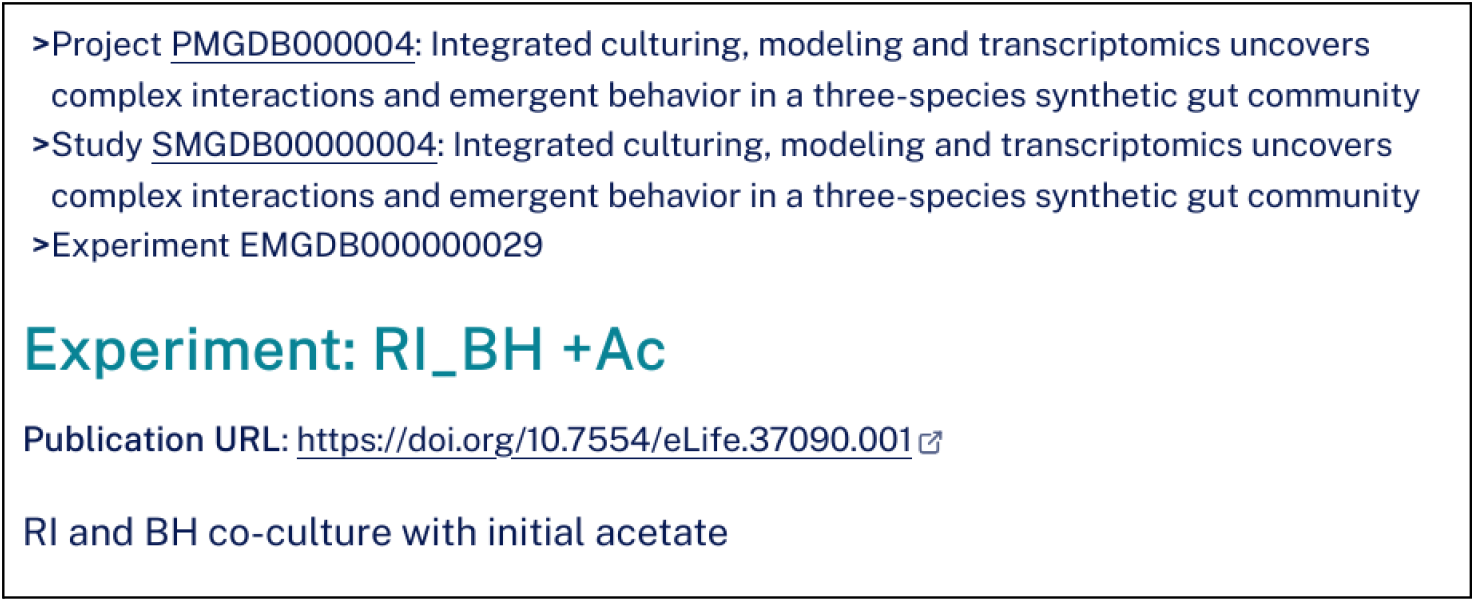
Illustration of an experiment record, its parent entities, and their unique identifiers

In addition to the *Communities* and the *Compartments*, the *Study* level also specifies measurement methods. Measurement methods describe measurement techniques, which fall into three main categories:

- Abundance measurements at the strain level. The following methodologies are currently supported: flow cytometry, plate counts, 16S rRNA sequencing, and qPCR.
- Abundance measurements at the community level, i.e., the total abundance of all community members. This is supported for the following methodologies: flow cytometry, optical density, and plate counts.
- Measurements of metabolite concentrations For additional details, a comprehensive overview of the main concepts underlying μGrowthDB is available under the “[Main concepts](https://mgrowthdb.gbiomed.kuleuven.be/help/main-concepts/)” tab on the Help page.

### Back-end and Front-end implementation

μGrowthDB’s backend is implemented in Python using the Flask web framework, with SQLAlchemy as the object-relational mapper, MySQL as the database management system, and the growthrates R package for growth curve fitting. The front-end makes use of JavaScript, specifically jQuery, and communicates with the server using AJAX. Plotly is used for the visualisation of growth curves. Figure 2 summarizes μGrowthDB’s design.

**Figure 2.**
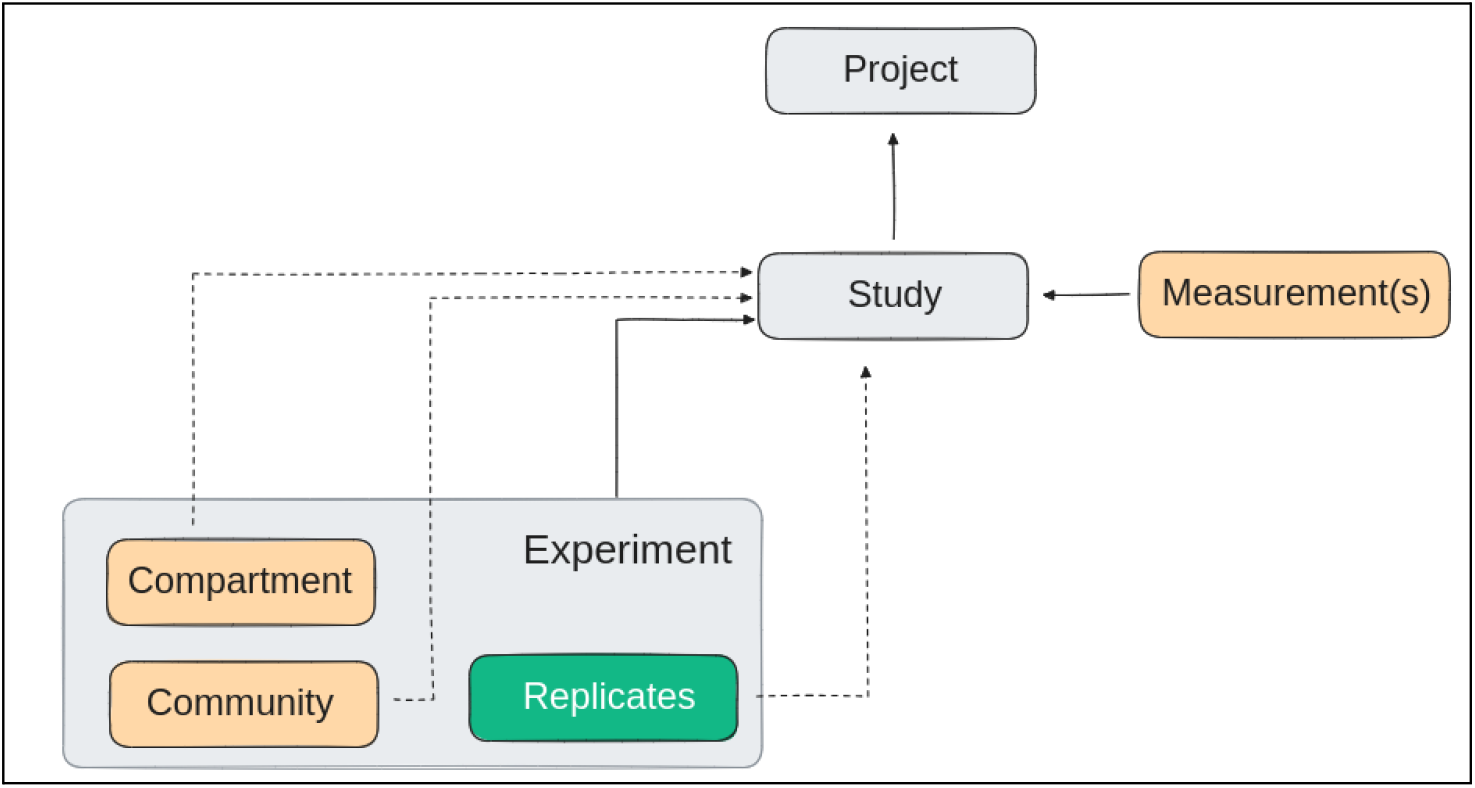
Key elements of the data scheme and backend-frontend design

The organization of the codebase follows a Model-View-Controller (MVC) architecture, similar to many popular web frameworks in use today, e.g. Rails (Bächle & Kirchberg, 2007), Django (*Django*, n.d.) and Symfony (Symfony, n.d.). The core logic is encapsulated in the classes and modules of the model layer. This makes it independent from any particular HTTP request, ensures its reusability for different use cases, and keeps it easily testable. The view layer holds HTML templates, their CSS styles, and JavaScript logic that adds interactivity to the user interface. The controller layer consists of HTTP request handlers that instantiate model code and inject its output into the appropriate view templates.

## Results

### Basic features in μGrowthDB

In μGrowthDB, users can query data sets by strain name or NCBI taxonomy ID, metabolite name or CheBI ID, study name or ID, and project name or ID via the *Search* page. Studies matching the query are listed with a table providing an overview of the techniques, strains, and metabolites involved. When a study is selected, more detailed information about the measurement techniques and an overview of its set of linked experiments is displayed, along with the URL to the publication if available. For each experiment, compartments with their properties, the strains involved, and perturbations (if any) are listed. Strains are linked to their NCBI taxonomy entry, media used in compartments are linked to their corresponding MediaDive entry if available, and metabolites are linked to their CheBI entry.

The user can either click the *Visualize* button on top of the screen to reach the study visualisation page or the plot icon (a red curve) next to an abundance or metabolite measurement technique to directly visualise the corresponding time series. In the visualisation page, the user can select which measurements to plot for which biological replicate(s) in which experiments. It is also possible to plot the mean across biological replicates, in which case standard deviations are displayed as well. In addition, the user can choose which time series to display on the left or right y-axis of the plot, which can represent different scales. This allows for an easy combination of metabolite and abundance measurements with their different measurement units. The user can also choose between a normal and a logarithmic y-axis and switch between units. The user-composed plot can be shared with a permalink obtained by clicking “Permalink to this page” below the plot.

Finally, the user can click the *Export data* button to download the study data or various sub-sets thereof in different formats. The help page provides information on database use, explanations of database concepts, and detailed help on data upload.

### Comparison of growth curves across studies

μGrowthDB enables users to compare growth curves across studies using the “shopping cart” principle. For this, the user can query studies for a strain or metabolite of interest, and in each *Study* select time series by clicking the *compare* icon (a double arrow) next to the biological replicate and measurement technique. The icon changes to a hand to indicate that the targeted measurement has been stored for comparison. The number of selected data sets is displayed next to the *Compare* menu item. Once all data sets of interest have been selected, the user can visit the *Compare* menu and display them together in a single plot, using the same plotting options as in the visualization page. The *Compare* menu also has a button to “empty the shopping cart”, i.e., to remove all data sets from the selection. The comparison workflow is illustrated in Figure 3. A permalink can be generated for the comparison plot, similar to the one available on the visualization page.

**Figure 3.**
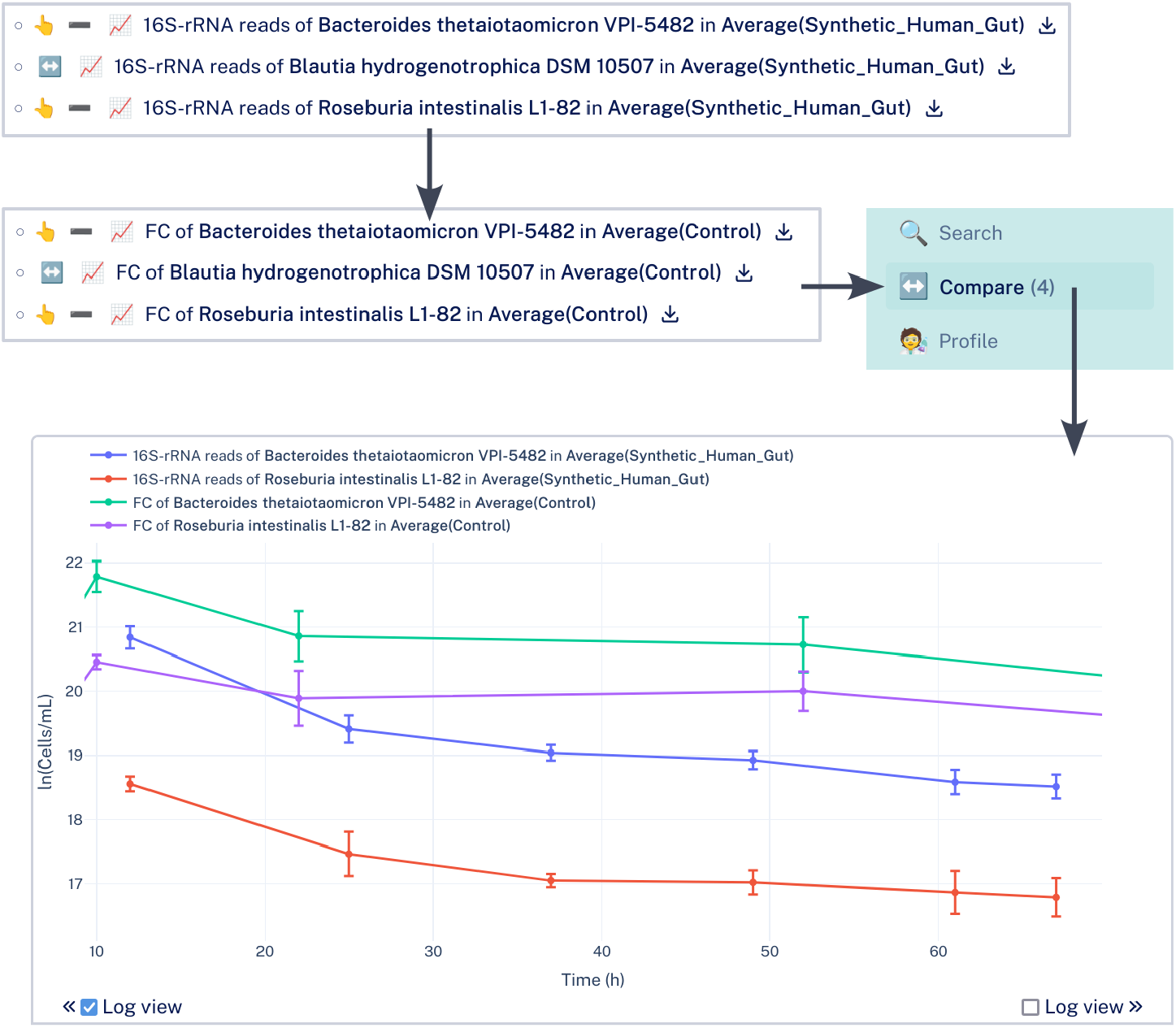
Example comparison between two strains measured in two different studies. In μGrowthDB, for each experiment, individual biological replicates as well as their average per strain and measurement technique (and *Compartment*) are provided. The upper left panels show experiments belonging to two different studies for which averages were selected for the same measurement technique for the two strains of interest, symbolized by the hand icon. The four selected data sets are collected in the *Compare* menu item shown on the right. The plot combining them is depicted in the bottom panel.

### Guided data upload

An important feature of μGrowthDB is the guided data upload. To upload or manage data, users need to log into their μGrowthDB account, which they can easily do with their ORCID on the *Log in* page; once logged in, this page is renamed to *Profile*. The upload process consists of seven well-documented steps supported by web forms. First, the user needs to describe the *Project* and *Study* concerned. It is possible to reuse a previous *Study* design for subsequent submission steps, so that microbial strains, measurement techniques, *Compartments*, and *Communities* do not have to be specified again. In the second step, the microbial strains involved in the *Study* need to be entered, which requires matching them to NCBI taxonomy entries. The form supports this step by automatically proposing NCBI taxonomy entries closest to the indicated strain name. Genetically modified strains can be entered as custom strains whose parent strain needs to match an NCBI taxonomy entry. In the third step, measurement techniques at the community and strain level need to be specified, and if any metabolites were measured, they need to be listed along with their measurement technique. Matching metabolites to ChEBI entries is supported in the same manner as for strains. In the fourth step, the *Compartments* used in the study need to be described, including their medium, atmosphere, inoculum, initial pH, and other properties. In the fifth step, the *Experiments* linked to the study need to be specified, i.e., which strains were grown in which compartments in how many biological replicates, and at which time points measurements were taken or perturbations applied. Based on the described experimental design, a study-specific data template is generated, which can then be downloaded by the user, filled out, and uploaded. Finally, in the seventh step, the user receives a link to the newly-created study that they can evaluate and revise. To protect sensitive data or allow for more time for checks, the study is only visible to the uploader until they click on the “Publish” button, making it available to the public. This is only possible after a period of 24 hours to encourage researchers to carefully investigate their upload. Once published, the data can then be queried and visualised as described above.

Basic data checks are performed during and after the submission, and user feedback is elicited to resolve non-unique names, confirm missing values, or deal with invalid values. The data upload process is summarised in Figure 4.

**Figure 4.**
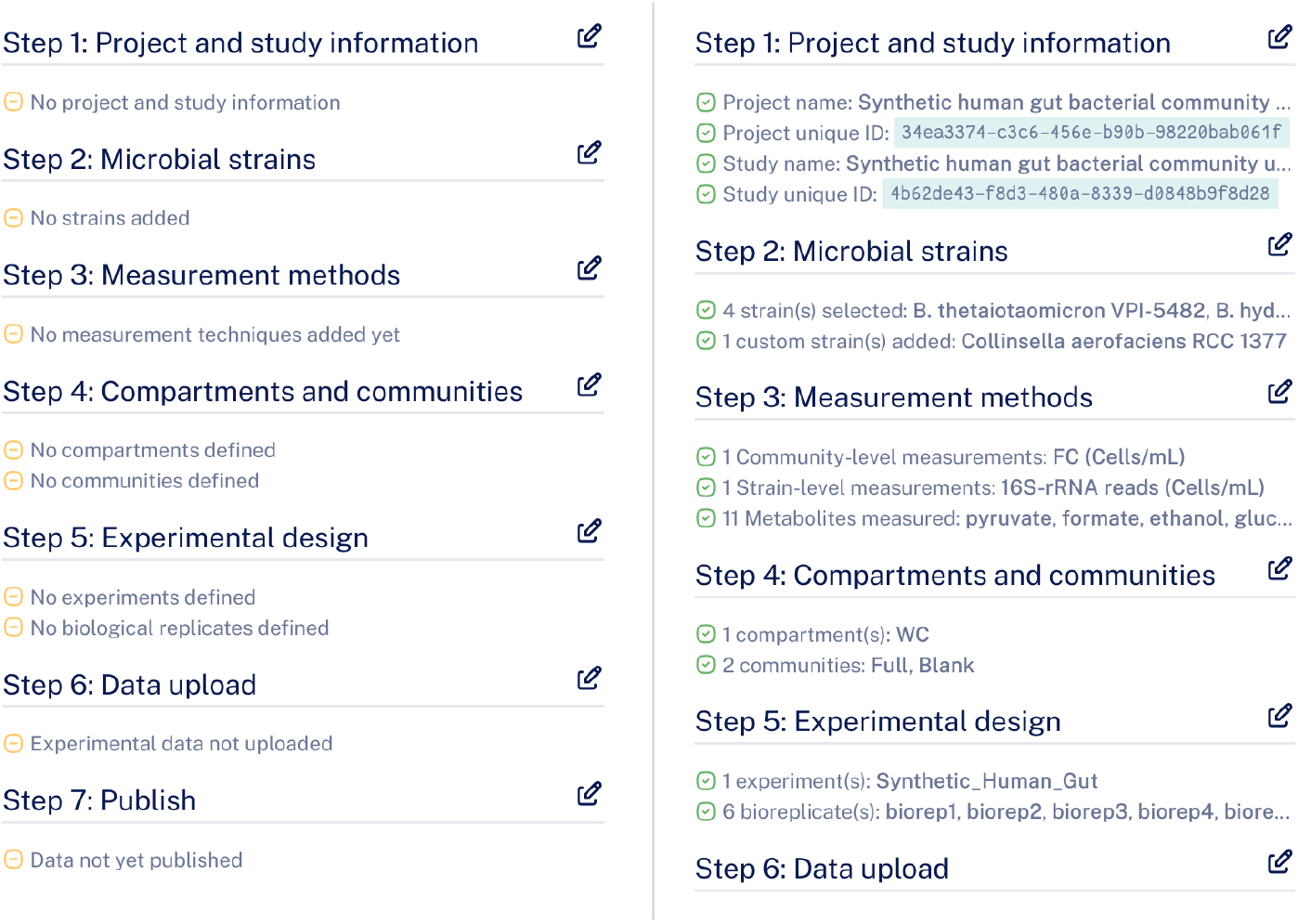
Summary of the upload process. Left: initial state of the form. Right: A filled-in form with short summaries of the entered study design.

The submitter of a study has access to a special button on the study page labeled “Manage”. This allows correcting study information and data if necessary. However, doing so implies repeating the upload process, resulting in a new version of the study. The old version is kept in the database, but queries will point to the latest version of a study.

### Growth curve fitting

For data submitters, basic growth curve fitting is available, including with the Easy-Linear method, which fits a line to the exponential phase in logarithmic space, and with the logistic and Baranyi-Roberts models (Baranyi et al., 1993; Baranyi & Roberts, 1994). The resulting fit is displayed together with the growth curve and the parameters; in particular, the growth rate, as well as the quality of the fit, are reported and can be downloaded as a CSV file. The growth curve fitting functionality is accessible via the *Manage* button.

### Local installation and code repository

μGrowthDB can be installed locally using micromamba as described in its GitHub repository: https://github.com/msysbio/bacterial_growth.

Feature requests and problems can be reported at μGrowthDB’s Discussions page: https://github.com/msysbio/bacterial_growth/discussions.

A guide for developers and further help is available at μGrowthDB’s ReadTheDocs page: https://mgrowthdb.readthedocs.io/.

### Future work & Call for community contribution

Developing a database for microbial growth curves involves several challenges; the biggest of which is populating the database. Data sets are recorded in a variety of formats scattered across the literature, and their submission cannot be automated. Thus, bulk upload is not possible, and a comprehensive number of data sets can only be attained through a community-wide effort. This is why we invested substantial work in making the data submission process as easy and flexible as possible. μGrowthDB success will strongly depend on the community’s continued efforts to populate it with high-quality data. We hope that this process, along with the incentives for data submission, such as growth rate calculation and visualization, will lead to a substantial contribution and thus, increase in the number of data sets in the near future.

We are currently developing a bulk download feature and an Application Programming Interface (API) to facilitate seamless access to the data. These additions will enable users to efficiently retrieve μGrowthDB datasets programmatically, further enhancing the resource’s accessibility and research utility.

We also plan to extend growth curve fitting to include more growth models, report confidence intervals of parameter values based on biological replicates, and to support fitting for every user (now only available for data submitters). In addition, we plan to develop a data submission checklist similar to those available for sequencing data (Yilmaz et al., 2011) to further ease the upload process. Ultimately, we hope that the submission of growth curve data to μGrowthDB will become standard when publishing such data in microbiology journals.

## Acknowledgements

We thank the microbiologists in the lab of microbial systems biology, including Charlotte van de Velde and Pallabita Saha, for their valuable feedback and Sebastian Proost for advice and beta testing. We also thank internship student Sourav Pattanaik for beta testing and data submission.

## Notes

https://mgrowthdb.gbiomed.kuleuven.be/

https://github.com/msysbio/bacterial_growth

### Competing Interest Statement

The authors have declared no competing interest.

